# Epididymal acquired sperm microRNAs modify post-fertilization embryonic gene expression

**DOI:** 10.1101/2023.10.01.560377

**Authors:** Natalie A. Trigg, Colin C. Conine

## Abstract

Sperm small RNAs have emerged as important non-genetic contributors to embryogenesis and offspring health. A subset of sperm small RNAs are thought to be acquired during epididymal transit. However, the transfer of RNAs from the somatic epididymis to sperm has been questioned, and the identity of the specific small RNAs transferred remains unclear. Here, we employ *Cre/Lox* genetics to generate germline- and epididymal-specific *Dgcr8* conditional knockout mice to investigate the dynamics of sperm microRNAs and their function in the early embryo. Testicular sperm from germline specific *Dgcr8* knockout mice have reduced levels of 98 microRNAs. Enthrallingly, following epididymal transit the abundance of 59% of these microRNAs are restored to control levels. Conversely, sperm from epididymal *Dgcr8* knockouts displayed a reduction of > 3.4-fold in 25 miRNAs. This substantial loss of epididymal miRNAs in sperm was accompanied by transcriptomic changes in the embryo which was rescued by microinjection of epididymal miRNAs. These findings ultimately demonstrate the acquisition of miRNAs from the soma by sperm during epididymal transit and their subsequent regulation of post-fertilization embryonic gene expression.

## INTRODUCTION

It is now well appreciated that, from a paternal perspective, the phenotype of healthy offspring is not solely dependent on the ability of sperm to simply fertilize the egg and deliver its haploid genome. Sperm also convey non-genetic information to the egg that have been implicated in embryonic development and the transmission of non-genetically inherited phenotypes (Champroux et al., 2018; Sharma, 2019). Among this non-genetic information are sperm small RNAs, for which several lines of evidence support their function as a form of paternal hereditary information (Kretschmer and Gapp, 2022; Zhang et al., 2019b). Indeed, many examples of paternal exposures ranging from stress to diet have demonstrated that the paternal environment can modulate the sperm small RNA payload to transmit non-genetically inherited phenotypes to progeny (Gapp et al., 2014; Rodgers et al., 2013; Sharma et al., 2018). A notable example is the phenocopying of offspring of stressed fathers in organisms developed from naïve zygotes microinjected with purified small RNAs from sperm of stressed males (Gapp et al., 2014; Rodgers et al., 2015; Wang et al., 2021). This causal link between exposure-induced sperm small RNA changes and the transmission of an offspring phenotype has also been demonstrated in the case of dietary perturbations in mice, such as Western-like diet fed fathers (Grandjean et al., 2015; Raad et al., 2021).

Aside from the role of sperm small RNAs under conditions of stress, the ‘physiologically normal’ pool of sperm small RNA delivered to the egg is involved in regulating early embryonic gene expression and development. The latter has been crudely demonstrated in mice using an approach in which sperm total RNA was introduced to a parthenogenetically activated egg (Conine et al., 2020). Diploid parthenotes develop efficiently to blastocysts and display broad transcriptomic changes throughout preimplantation embryonic development compared to sperm-fertilized embryos (Conine et al., 2020; Park et al., 2013). Fascinatingly, the introduction of sperm RNA to parthenotes results in changes in expression of hundreds of genes to more closely resemble the expression of a sperm fertilized embryo (Conine et al., 2020). In narrowing the role of individual small RNA classes in regulating post-fertilization gene expression, several additional studies have specifically identified post-fertilization functions for sperm microRNAs (miRNAs) in the early embryo (Wang et al., 2022; Yuan et al., 2016). Embryos produced via intracytoplasmic sperm injection (ICSI) of sperm conditionally depleted of miRNAs during spermatogenesis display reduced preimplantation development which was partially rescued by microinjection of sperm small RNA (Yuan et al., 2016). These findings specifically highlight a function of sperm miRNAs post-fertilization in the early embryo that we are only beginning to understand.

Following sperm development in the testis, spermatozoa are passaged to the epididymis, where they undergo extensive molecular changes that culminate in a mature, motile, functionally competent sperm cell (Gervasi and Visconti, 2017). Among the alterations that occur during this process is the modulation of the sperm small RNA profile. While sperm leaving the testis harbor a small RNA profile primarily comprised of PIWI interacting RNAs (piRNAs), the small RNA profile of mature cauda spermatozoa is dominated by transfer-RNA derived fragments (tRFs) (Hutcheon et al., 2017; Sharma et al., 2018). While the population of miRNAs within sperm proportionally decreases along the epididymis, a subset of miRNAs are enriched in cauda sperm compared to sperm isolated from the proximal (caput) epididymis (Belleannée et al., 2012; Conine et al., 2018; Nixon et al., 2015). This dynamic post-testicular expression is thought to occur on ∼100 miRNAs, highlighted by miRNAs belonging to the miR-34b/c and miR-17-92 family, and those of the miR-880 X-chromosome cluster. Interestingly, these miRNAs are also detected in testicular sperm and therefore are seemingly lost upon entry to the epididymis, before being reacquired (Conine et al., 2018). Owing to the transcriptionally and translationally inert state of epididymal sperm (Johnson et al., 2011), it is likely that such changes to the sperm small RNA payload is driven by external factors. Among the mechanisms thought to facilitate the dynamics of sperm small RNAs are extracellular vesicles produced by the epididymal epithelium, termed epididymosomes. These vesicles harbor a rich profile of small RNA, including miRNAs and tRFs (Reilly et al., 2016; Sharma et al., 2016). Further, *in vitro* co-incubation experiments and *in vivo* tracking of metabolically labeled RNAs have demonstrated the transfer of miRNAs from the epididymis to sperm, via epididymosomes (Chan et al., 2020; Reilly et al., 2016; Sharma et al., 2018). Despite this evidence, a recent study has suggested an alternative mechanism underlying the dynamics of sperm small RNAs in the epididymis which occurs through selective shuffling of RNA between sperm and remnant germ cell cytoplasm that remains adhered to the midpiece of the sperm cell, known as the cytoplasmic droplet, rather than small RNA acquisition from the soma (Wang et al., 2023). While this study provides compelling evidence to support this mechanism in the transfer of tRFs and rRNA-derived RNA fragments (rRFs), it does not explain the post-testicular dynamics of changes in sperm miRNAs in the epididymis as miRNAs are scarcely abundant in sperm cytoplasmic droplets (Wang et al., 2023). Thus, these findings leave open the possibility that miRNAs are indeed transferred from the soma to sperm during epididymal transit. To address this, we exploited a genetic approach to conclusively demonstrate that sperm acquire miRNAs from the epididymis and that these epididymal miRNAs function post-fertilization in the early embryo.

## MATERIALS AND METHODS

### Ethics Statement

All animal care and use procedures were in accordance with the guidelines of the Children’s Hospital of Philadelphia and University of Pennsylvania Institutional Animal Care and Use Committee (CHOP IACUC Protocol #: IAC 23-001364, UPenn PSOM IACUC Protocol #: 806911).

### Experimental animals

To achieve selective genetic inactivation of DiGeorge syndrome critical region 8 (*Dgcr8*) and hence disrupt the miRNA pathway in the male reproductive tract, we bred *Dgcr8*^flox^ (Dgcr8^fl/fl^) female mice with male mice expressing *iCre* under the Stimulated by retinoic acid 8 (Stra8, JAX stock #008208) or Defensin beta 41 (Defb41) promotors, for testis specific or epididymal specific knockout, respectively (Björkgren et al., 2012; Sadate-Ngatchou et al., 2008). The *Dgcr8* floxed allele contains loxP sites flanking exon 3, hence cross breeding with *Cre* strains causes the deletion of exon 3 that leads to a frameshift and multiple premature stop codons (MMRRC stock #32051) (Rao et al., 2009). All animals were genotyped by classic PCR as described previously (Sharma et al., 2018). For assisted reproductive techniques (IVF and ICSI) and breeding trials to assess fertility in conditional knockout males, 5–7-week-old female FVB/NJ mice were used as oocyte donors.

### Histology, SDS-PAGE and immunoblotting

Testis and epididymal tissue from experimental mice were fixed in 10% buffered formalin for 48 h before washing in 50% ethanol. Fixed tissue was embedded in paraffin and 5 μm sections were stained with hematoxylin and eosin (H&E) at the University of Pennsylvania School of Veterinary Medicine Comparative Pathology Core and imaged on Aperio VERSA 200 platform (Leica). Additionally, testis and epididymal tissue were homogenized in RIPA buffer on ice for 15 min, centrifuged at high speed (15 min, 13,000 × *g*) and protein containing supernatant collected for immunoblotting to examine *Dgcr8* protein expression. Proteins were separated by SDS-PAGE and gels were transferred to nitrocellulose membranes and blocked for 1 h in 3% BSA / TBS supplemented with 1% Tween-20 (TBST). After washing the membrane in TBST, anti-DGCR8 (Abcam, Cat# ab191875) antibodies diluted (1:1000) in 1% BSA / TBST were added to the membrane and incubated overnight at 4°C. Following three TBST washes, the membrane was incubated in secondary antibodies for 1 h at room temperature before washing three times. Proteins were detected using enhanced chemiluminescence reagents as per the manufacturer’s instructions (Cytiva, Cat# RPN2109).

### Testicular sperm collection

Testicular sperm was obtained by removing the tunica albuginea and mincing testes in 0.15 M NaCl to release spermatozoa into the media. Sperm were allowed to disperse for 10 min and then the tissue suspension was transferred to a new tube and incubated on ice to allow tissue to settle. The top 1 mL was loaded onto a 52% isotonic Percoll solution and centrifuged at 27,000 × *g* for 10 min at 10°C (Optima-XPN-80, Beckman Coulter). Following centrifugation, the supernatant was removed, leaving the pellet and ∼1 mL solution, which was subsequently washed in 10 mL 0.15 M NaCl. The sperm pellet was then subjected to three washes in 0.75 M NaCl before incubation in lysis buffer (0.1% SDS and 0.5% Triton-X 100) to remove contaminating somatic cells. Cells were washed in PBS following confirmation of somatic cell elimination and the sperm pellet was snap frozen for RNA extraction.

### Epididymal sperm collection and assessment of sperm parameters

To isolate mature epididymal sperm, dissected cauda epididymides were placed in a dish with Whitten’s media (100 mM NaCl, 4.7 mM KCl, 1.2 mM KH_2_PO_4_, 1.2 mM MgSO_4_, 5.5 mM glucose, 1 mM pyruvic acid, 4.8 mM lactic acid (hemicalcium), and HEPES 20 mM) and sperm were retrieved by making an incision in the epididymis and gently squeezing luminal contents into the media. Following a ‘swim out’ period of 10 min sperm suspensions were transferred to a tube allowing the spermatozoa to ‘swim up’ for an additional 5 min at 37°C. The supernatant was retrieved, and sperm cells were pelleted by centrifugation at 10,000 × *g* for 5 mins. Sperm pellets were washed in PBS before incubation in somatic cell lysis buffer for 10 min on ice to remove contaminating somatic cells. Samples were assessed for contamination using phase microscopy. Purified sperm were then washed in PBS, snap frozen in liquid nitrogen and stored at -80°C ready for RNA extraction.

Sperm parameters were assessed following the initial ‘swim up’ incubation. Sperm motility and morphology was assessed using phase contrast microscopy. Trypan blue vitality stain was mixed 1:1 with sperm solution to determine sperm viability via the exclusion of stain in live cells. For each of these parameters a minimum of 100 cells were counted.

### Isolation of epididymal extracellular vesicles

Epididymosomes were isolated as previously described (Sharma et al., 2016). Briefly, epididymal luminal contents devoid of spermatozoa were centrifuged at 10,000 × *g* for 5 min, then 30 min at 4°C to remove all cellular debris. The resulting supernatant was subjected to ultracentrifugation at 120,000 × *g* at 4°C for 2 h (Optima Max TL, Beckman Coulter). Pellets were resuspended in 1 ml cold PBS and centrifuged at 120,000 × *g* at 4°C for 2 h. The epididymosome pellet was resuspended in 30 µL cold PBS and snap frozen ready for downstream RNA extraction.

### Pregnancy outcomes analysis

The reproductive capacity of heterozygous and homozygous *Dgcr8^fl/fl^; Defb41-iCre* males was assessed by mating a male with one control female and checking for vaginal plugs daily to confirm mating (n = 5-6 individual males in the breeding scheme). Once mating was confirmed, plugged females were separated from the male and sacrificed on day 17.5 post-copulation for pregnancy outcomes. Intact uteri were dissected, and the total number of viable and resorbing implantations were recorded.

### RNA extraction

To extract RNA from populations of sperm, samples were lysed in sperm lysis buffer (6.4 M Guanidine HCl, 5% Tween 20, 5% Triton, 120 mM EDTA, and 120 mM Tris pH 8.0) with 0.66 mg/mL Proteinase K and 33 mM DTT and incubated at 60°C for 15 min with shaking. One volume of water and 2 volumes of TRI reagent was added and mixed thoroughly to ensure complete lysis. The samples were transferred to phase lock tubes (Quantabio, Cat# 2302830) and 0.2 × volume of BCP (1-bromo-2 chloropropane) was added. To mix, tubes were inverted 15 times, and centrifuged at 14,000 × *g* for 4 mins at 4°C. The aqueous phase was transferred to a fresh low bind tube before adding 20 µg glycoblue and 1.1 × volume of isopropanol. Samples were then mixed well, and the RNA allowed to precipitate for at least 1h at -20°C. Following precipitation, RNA was pelleted at 14, 000 × *g* for 15 min at 4°C, then washed in 70% ice cold ethanol and reconstituted in water. For tissue RNA extraction, tissue was homogenized in TRI reagent until complete cell breakdown. The samples were transferred to phase lock tubes with BCP before phase separation and isopropanol precipitation as above.

### Small RNA-sequencing

The small RNA fraction (18-40 nucleotides) was purified from total RNA samples using 15% polyacrylamide-7M urea denaturing TBE gel electrophoresis. Size selection of small RNA was followed by ligation-dependent small RNA library construction, as previously outlined, using a modified Illumina Tru-Seq small RNA library prep protocol (Sharma et al., 2016). Briefly, the 3′ ends of small RNA were ligated to an adapter using T4 truncated RNA ligase (Lucigen, Cat# LR2D11310K and NEB, Cat# M0242L), followed by the ligation of the 5′-end adaptors by T4 ligase. Ligated RNAs were then converted to cDNA using Superscript II. To control for differences in the amount of starting material, samples representing the range of input were selected for amplification optimization to determine the minimum number of cycles without overamplifying. Once determined, DNA was amplified by sequential rounds of PCR and individual libraries were subsequently gel purified to remove empty adapters and nonspecific amplicons. Libraries were quantified and pooled for sequencing on a NextSeq 1000. Read quality was assessed using FastQC, and adapter sequences were trimmed using trimmomatic. Trimmed reads were mapped sequentially to rRNA mapping reads, miRbase, murine tRNAs, pachytene piRNA clusters (Li et al., 2013), repeatmasker and Refseq using Bowtie 2 and totaled using Feature counts (Yukselen et al., 2020). Mapped reads were normalized to parts per million (total genome mapping reads) for subsequent analysis. Raw read count data (Table S1-2) was normalized and analyzed for differentially abundant small RNAs using DESeq2 (Love et al., 2014). Differentially abundant small RNAs were determined as those with a fold-change > 2 and adjusted *P*-value < 0.05.

### Intracytoplasmic sperm injection (ICSI) and microinjection

Sperm were prepared for intracytoplasmic sperm injection (ICSI) by washing pelleted sperm in PBS and sonicating the sperm suspension to detach sperm heads from tails for 5 sec at low power. Following sonication, sperm were washed and resuspended in modified nuclear isolation media (NIM) with 0.1% polyvinyl alcohol (PVA) ready for sperm injection. Oocytes were obtained from superovulated female mice, dissociated from cumulus cells by incubation in hyaluronidase (3 mg/mL), washed in KSOM, and incubated in KSOM at 37°C, 5% CO_2,_ 5% O_2_ until microinjection.

An aliquot of sperm was aspirated into a droplet of NIM 0.1% PVA on the injection plate and eggs were transferred for injection into droplets of FHM (Millipore, Cat# MR-122-D) with 1% PVA in groups of 10-15. The injection needle (Eppendorf, Cat# 930001091) was washed in NIM 0.1% PVA proceeding and between injections. Following sperm injection, presumptive zygotes were washed through 6 drops of KSOM (Millippore, Cat# MR-101-D) and cultured in KSOM at 37°C, 5% CO_2,_ 5% O_2_. Zygotes to be microinjected were first allowed to incubate for 1 h following ICSI before microinjection. Zygotes were then transferred to the injection plate, in droplets of FHM 1% PVA for micromanipulation. Control injections consisted of H3.3-GFP mRNA alone (30 ng/µL) and experimental injections included H3.3-GFP mRNA in combination with purified miRNAs from cauda epididymosomes (2 ng/µL). RNA injections were carried out using a Femtojet (Eppendorf) microinjector with Femtotip II microinjection capillary tips (Eppendorf, Cat# 930000043) at 100 hPa pressure for 0.2 seconds, with 7 hPa compensation pressure. After RNA injection, zygotes were washed in KSOM and continued to be cultured. The presence of H3.3-GFP was confirm using fluorescence microscopy and GFP-positive 2-cell embryos were separated, cultured, and collected for single-embryo mRNA-sequencing. Four-cell and morula stage embryos were collected into 1 × TCL buffer (Qiagen, Cat# 1070498) containing 1% 2-mercaptoethanol at 46 and 72 h post-fertilization, respectively.

### Single-embryo mRNA-sequencing

Single-embryo mRNA-seq libraries were constructed using the SMART-Seq protocol (Conine et al., 2018; Ramsköld et al., 2020; Sharma et al., 2016; Trombetta et al., 2014). Briefly, after embryo cell lysis, RNA was isolated via RNAClean-XP beads (Beckman Coulter, Cat# A63987) and full-length polyadenylated RNA was reverse transcribed using Superscript II (Invitrogen, Cat# 18064014). The cDNA was then amplified using 10 cycles and subsequently (0.33 ng) used to construct a pool of uniquely indexed samples using the Nextera XT kit (Illumina, Cat# FC-131-1096). Finally, pooled libraries were sequenced on a NextSeq 1000. Data were mapped against the *Mus musculus* genome (mm10) using RSEM and normalized to transcripts per million (tpm) (Table S4-6). Single embryos with less than 6,000 (4-cell) or 8,000 (morula) genes with expression greater than 1 tpm were removed from analysis. To assess the differentially expressed genes, data was loaded into R Statistical Software and analyzed using the DESeq2 package using a negative binomial distribution (Love et al., 2014). For all single-embryo mRNA-seq experiments differentially expressed genes (DEGs) were determined by a fold-change > 1.5 and *P*-value < 0.01. This less stringent threshold was selected to allow for the comparison of a larger pool of genes across all embryo experiments and analysis using IPA software. Nevertheless, the same conclusions are drawn from our data when multiple hypothesis testing (adjusted *P*-value) is employed.

### Statistical analysis

GraphPad Prism (version 9.5.1) was used for statistical analysis. Normality of datasets was assessed with the Shapiro-Wilk test (α = 0.05). Following this, to assess the difference of two means an unpaired Student’s t-test was used for normally distributed data and the non-parametric Man-Whitney U test was performed for data where normality failed. To assess differences between development of control and knockout sperm fertilized embryos, one-way ANOVA with Sidak multiple comparison test was used. Data is presented as mean values ± standard error of the mean.

## RESULTS

### Generation of GC-*Dgcr8* and Epi-*Dgcr8* mice

The *Dgcr8* gene encodes a double-stranded RNA binding protein which forms a subunit of the microprocessor complex. This complex facilitates the processing of the primary miRNA transcript, hence the loss of *Dgcr8* directly impacts miRNA biogenesis (Landthaler et al., 2004). Testis and epididymis specific genetic inactivation of *Dgcr8* in the male reproductive tract was achieved by breeding *Dgcr8*^flox^ (Dgcr8^fl/fl^) female mice with either *Stra8*- or *Defb41*-*iCre* male mice, respectively. *Stra8* is initially expressed in the differentiating spermatogonia at postnatal day 3 (Oulad-Abdelghani et al., 1996), achieving a germ cell specific knockout of *Dgcr8*. Conversely, *Defb41* is expressed in the epithelium of the epididymis, providing an epididymal specific knockout (Jalkanen et al., 2005). For simplicity, germ cell specific ablation of *Dgcr8* (*Dgcr8^fl/fl^; Stra8-iCre)* will be referred to as GC-*Dgcr8*, while the epididymal conditional knockout (*Dgcr8^fl/fl^; Defb41-iCre)* will be referred to as Epi-*Dgcr8* (Fig. 1A). Immunoblot analysis revealed successful reduction of *Dgcr8* protein expression in the testis and caput epididymis, in GC-*Dgcr8* and Epi-*Dgcr8* knockout mice, respectively (Fig. S1A-B).

**Figure 1:**
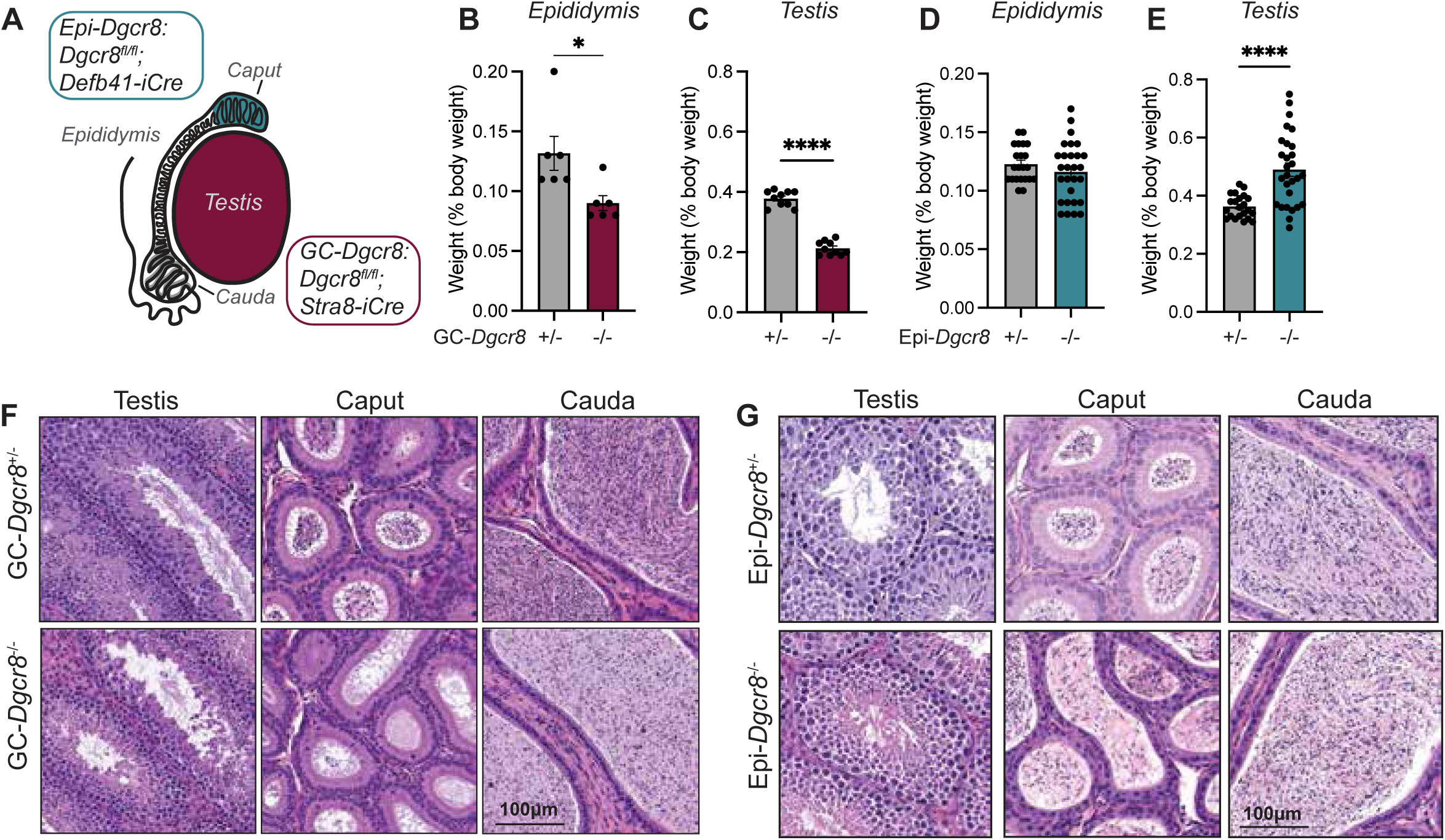
Loss of *Dgcr8* in germ cells and the epididymis impacts the male reproductive tract. (A) Schematic of i*Cre* driven conditional knockout of *Dgcr8* in the testis and the epididymis. (B-E) The testes and epididymides of 8–12-week-old male mice were dissected and weighed for GC-*Dgcr8* (B-C) and Epi-*Dgcr8* (D-E) heterozygous controls (+/-) and sibling matched homozygous mutant males (-/-). Tissue weights are reported as a percentage of mouse body weight. (F-G) Testes and epididymides from GC-*Dgcr8^-/-^* (F) and Epi-*Dgcr8^-/-^* (G) male mice and their respective controls were fixed in 10% buffered formalin and H&E stained to assess morphology. Data is represented as mean ± SEM with each dot representing a measurement from a single mouse or testis/epididymis. Statistical significance is denoted as *P* ≤ 0.05 where asterisks represent levels of significance. **** = *P* < 0.0001.

### *Dgcr8* conditional knockout influences testes weight and sperm functionality

After sacrificing adult male mice (8–12-week-old), we noted no difference in mouse weight between control and *Dgcr8^-/-^* males, despite the site of conditional knockout (both GC and Epi; Fig. S1C). However, the loss of *Dgcr8* impacted testicular and epididymal weight. GC-*Dgcr8*^-/-^ males displayed reduced testicular and epididymal to body weight ratios compared to control males, while Epi-*Dgcr8^-/-^* males exhibited increased testicular to body weight ratios and no change in epididymal to body weight ratio (Fig. 1B-E, S1C-D). Macroscopic evaluation of 2-month-old GC-*Dgcr8*^-/-^ testes revealed impaired spermatogenesis, at the late spermatid stage, with a reduced number of spermatocytes and testicular sperm identified. Consequently, epididymal sections of GC-*Dgcr8*^-/-^ mice revealed epididymal tubules with reduced sperm content (Fig. 1F). Conversely, Epi-*Dgcr8^-/-^* male mice displayed normal spermatogenesis but revealed an underdeveloped initial segment of the epididymis that was difficult to distinguish from the caput epididymis as has been previously reported in epididymal conditional knockouts of *Dicer1* (Fig. 1G, S1E) (Björkgren et al., 2012).

Scoring sperm parameters of populations of cauda sperm from GC-*Dgcr8*^-/-^ males revealed reduced sperm motility in GC-*Dgcr8*^-/-^ sperm compared to control sperm, but no accompanying loss of vitality (Fig. 2A-B). Accordingly, GC-*Dgcr8*^-/-^ males sired litters with a reduced number of pups compared to control males (Fig. 2C). In agreement with natural mating results, fertilization rates from IVF with GC-*Dgcr8*^-/-^ cauda sperm was significantly reduced compared to control sperm fertilized embryos (Fig. 2D). However, using ICSI, GC-*Dgcr8*^-/-^ cauda sperm fertilized embryos at rates similar to heterozygous (GC-*Dgcr8*^+/-^) sperm, a result that was mirrored using testicular sperm (Fig. S2A-B). Assessment of Epi-*Dgcr8*^-/-^ sperm parameters revealed a significant decrease in sperm motility (Fig. 2E), which displayed a precipitous decline in male mice 10 weeks or older (Fig. 2F). Reduced sperm motility was accompanied by a modest but significant reduction in sperm vitality (Fig. 2G). Subsequently, Epi-*Dgcr8*^-/-^ males consistently failed to produce pups via natural mating despite males readily mating with females, which was confirmed via the presence of a copulatory plug (Fig. 2H). Further, Epi-*Dgcr8*^-/-^ sperm failed to fertilize embryos with IVF, even when young adult (8-10-week-old) male mice were utilized (Fig. S2C). This failed IVF fertilization was initially attributed to reduced sperm motility, which was further supported by the assessment of sperm motility over time *in vitro* which revealed a steady decline in Epi-*Dgcr8*^-/-^ sperm motility over 60 min (Fig. S2D). To overcome this sperm motility defect, we used ICSI with Epi-*Dgcr8*^-/-^ sperm to fertilize eggs. Using ICSI, Epi-*Dgcr8*^-/-^ sperm were able to fertilize control eggs at levels comparable to control sperm (Fig. 2I). However, scoring embryo development over 96 h revealed a reduced number of embryos that were fertilized by Epi-*Dgcr8*^-/-^ sperm developing to the blastocyst stage, compared to control sperm fertilized embryos. Only 35% of Epi-*Dgcr8*^-/-^ sperm injected eggs developed to the blastocyst stage compared to nearly 60% in control sperm injected embryos. Indeed, Epi-*Dgcr8*^-/-^ sperm injected eggs arrested at 1-cell (21% vs 12% in controls) and 2-cell (19.5% vs 6%, *P* < 0.05) stages of development (Fig. 2J-K).

**Figure 2:**
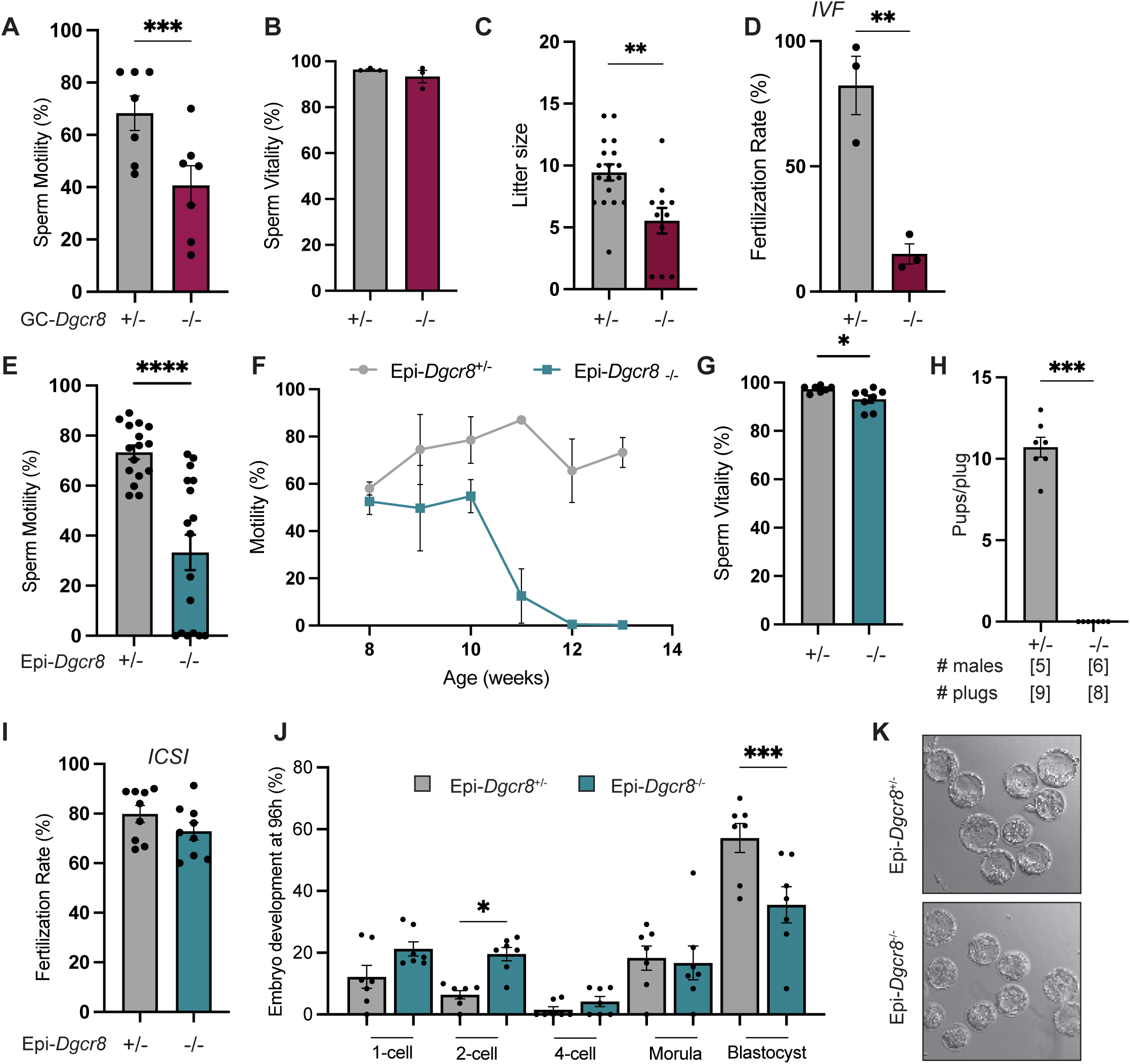
Sperm functionality and fertility is impacted in conditional *Dgcr8* knockout mice. (A-B) Populations of control and GC-*Dgcr8*^-/-^ cauda spermatozoa were assessed for sperm motility (A) and vitality (B) using phase contrast microscopy. (C) Average number of pups born following natural mating of control and GC-*Dgcr8*^-/-^ male mice. (D) Fertilization rate of GC-*Dgcr8*^-/-^ cauda sperm via *in vitro* fertilization (IVF). (E) Average sperm total motility of control and Epi-*Dgcr8*^-/-^ cauda sperm and (F) plotted as a function of mouse age. (G) Sperm vitality of Epi-*Dgcr8*^-/-^ and control cauda sperm. (H) Average number of pups born per plug for control and Epi-*Dgcr8*^-/-^ males. (I) Fertilization rate and (J) 96 h development of embryos fertilized by Epi-*Dgcr8*^-/-^ cauda sperm via intracytoplasmic sperm injection (ICSI) compared to control sperm fertilized embryos. (K) Representative phase images at 96 h of embryos fertilized by control or Epi-*Dgcr8*^-/-^ sperm. Data is presented as mean ± SEM with each dot representing a measurement from a single mouse. Statistical significance is denoted as *P* ≤ 0.05 where asterisks represent levels of significance. * = *P* < 0.05, ** = *P* < 0.01, *** = *P* < 0.001 and **** = *P* < 0.0001.

### Epididymal transit modifies the sperm microRNA payload

To determine how ablation of the miRNA pathway in the developing germ cells or the epididymis impacts the sperm miRNA profile, we performed small RNA-seq on testicular and cauda (mature) sperm from males with the miRNA pathway knocked out in the testis and epididymis (Fig. S2G). Expectedly, the depletion of *Dgcr8* in early germ cells led to testicular sperm with aberrant miRNA levels compared to controls. Namely, the abundance of 105 miRNAs were significantly altered in GC-*Dgcr8*^-/-^ sperm (Fig. 3A). Importantly, the majority of these altered miRNAs (93%) were decreased in abundance compared to control testicular sperm. Interestingly, however, once sperm had traversed the epididymis, the number of miRNAs displaying significantly altered abundance was reduced to 69 miRNAs (Fig. 3C). Indeed, all the miRNAs that displayed reduced abundance in GC-*Dgcr8*^-/-^ testicular sperm, demonstrated fold changes with less magnitude in cauda sperm, with 59% of the miRNAs altered in testicular sperm no longer significantly differentially expressed in GC-*Dgcr8*^-/-^ cauda sperm compared to control sperm (58 of the 98 reduced miRNAs in GC-*Dgcr8*^-/-^ testis sperm). Conversely, testicular sperm from Epi-*Dgcr8^-/-^* mice harbored normal levels of miRNAs, with only a few differentially altered miRNAs identified between Epi-*Dgcr8^-/-^* and control sperm (Fig. 3B). However, depletion of *Dgcr8* in the epididymal epithelium led to an altered miRNA profile in sperm isolated from the cauda epididymis (Fig. 3D). Specifically, 55 miRNAs displayed altered abundance, of which, the majority (30 miRNAs) were increased in abundance in Epi-*Dgcr8*^-/-^ sperm. Nevertheless, the greatest magnitude of change (up to 9-fold reduction) was seen in the 25 miRNAs that were significantly reduced in Epi-*Dgcr8*^-/-^ sperm compared to control sperm, with an average fold reduction of 3.4-fold among these 25 miRNAs. Notably, 21 of the significantly reduced miRNAs have been previously identified to be specifically enriched in cauda compared to caput (early epididymal) sperm, including miRNAs from the miR-880 X-chromosome cluster and miR-34/449 family (Conine et al., 2018; Nixon et al., 2015). Further, 16 of the 25 reduced miRNAs were also reduced in GC-*Dgcr8*^-/-^ testicular sperm, but to a lesser extent (Fig. 3E). Importantly, the 16 miRNAs that were decreased in GC-*Dgcr8*^-/-^ testicular sperm and Epi-*Dgcr8*^-/-^ cauda sperm, compared to control sperm were subsequently increased in GC-*Dgcr8*^-/-^ cauda sperm, highlighting the acquisition of these miRNAs during epididymal transit (Fig. 3E).

**Figure 3:**
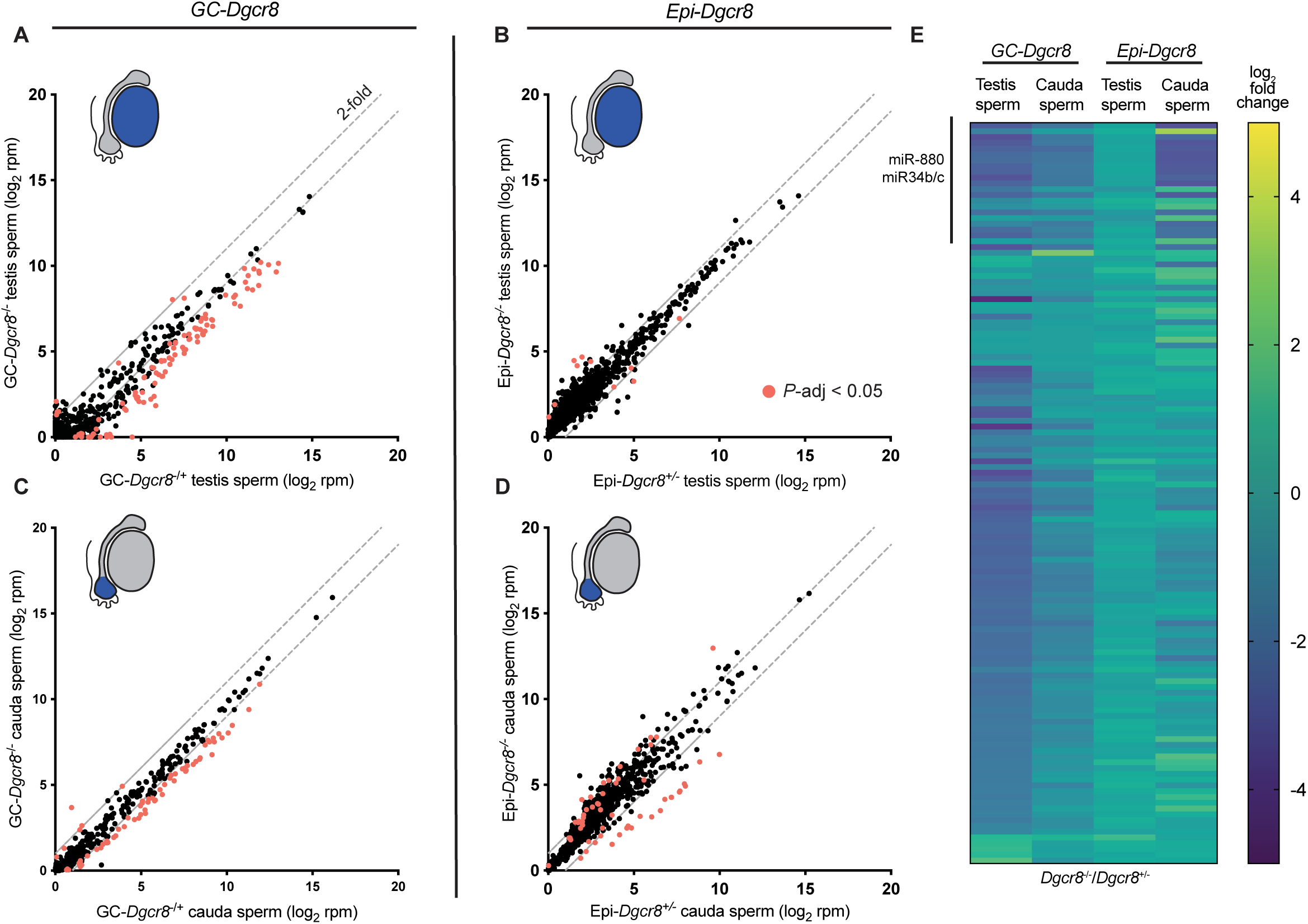
Epididymal transit modifies the sperm microRNA payload. Populations of testicular and cauda spermatozoa from control (heterozygous siblings) and mutant GC-*Dgcr8* (left) and Epi-*Dgcr8* (right) mice were prepared for small-RNA sequencing (n= 4-8). (A-D) Scatter plots depict miRNA abundance (reads per million genome mapping reads; rpm) for *Dgcr8*^+/-^ (x-axis, log_2_) sperm versus *Dgcr8*^-/-^ (y-axis, log_2_) sperm. miRNA abundance in: (A) GC-*Dgcr8* and (B) Epi-*Dgcr8* small RNA-seq from testicular sperm, and (C) GC-*Dgcr8* and (D) Epi-*Dgcr8* cauda spermatozoa. Orange dots represent statistically significant differentially accumulated miRNAs (*P*-adj ≤ 0.05) in *Dgcr8*^-/-^ sperm compared to control sperm. Blue shading in testis and epididymis schematic indicates origin of sperm population. (E) Heatmap depicting the fold change of miRNAs differentially accumulated (adjusted *P*-value ≤ 0.05 and fold-change ≥ 2) in either GC-*Dgcr8*^-/-^ testicular sperm and/or Epi-*Dgcr8*^-/-^ cauda sperm. Data is presented as log_2_ fold change of *Dgcr8^-/-^* sperm compared to *Dgcr8^+/-^* sperm.

### Epididymal miRNA and mRNA expression is altered in *Dgcr8* conditional knockout mice

Small RNA sequencing of caput epididymal tissue revealed large disruptions in miRNA levels in Epi-*Dgcr8*^-/-^ mice compared to control mice (Fig. 4A). Further, in examining miRNA abundance in the cauda epididymis of Epi-*Dgcr8*^-/-^ mice we observed a significant reduction in 18 miRNAs, including 12 that were also reduced in populations of Epi-*Dgcr8*^-/-^ cauda sperm (Fig. 4B). This finding suggests that epididymal miRNA levels within one epididymal segment can influence small RNA expression in subsequent regions, a form of paracrine signaling that has been previously demonstrated for miRNA-mRNA regulation in the epididymis (Belleannée et al., 2013; Belleannée et al., 2012). Thus, next we examined mRNA changes in epididymal tissue samples. This analysis identified that the loss of *Dgcr8* in the caput epididymis disrupted the expression of 981 genes, of which approximately half were up-regulated (497 genes) and the remaining were down-regulated (484 genes) (Fig. 4C). In contrast, only modest transcriptomic changes were evident in the cauda epididymis, where 32 genes were significantly altered compared to control samples (Fig. 4D). Comparatively, mice lacking *Dgcr8* in testicular germ cells demonstrated epididymal miRNA levels largely comparable to control epididymides (Fig. S3A-B). Indeed, no altered miRNAs were identified in the caput epididymis and 6 miRNAs were significantly reduced in GC-*Dgcr8*^-/-^ cauda epididymal tissue compared to controls. Further, mRNA-sequencing of epididymal tissue of GC-*Dgcr8* males identified the up-regulation of 80 genes and down-regulation of 3 genes in the caput epididymis. In the cauda epididymis, only 6 genes were altered, displaying increased expression in GC-*Dgcr8*^-/-^ cauda epididymis compared to controls (Fig. S3 C-D). Ingenuity Pathway Analysis (IPA) software identified several male fertility related pathways inhibited in the epididymis of GC-*Dgcr8*^-/-^ males (Fig. 4E). Interrogation of this analysis also identified predicted upstream regulators of the transcriptomic changes in *Dgcr8*^-/-^ epididymides, including the predicted activation of interferon-gamma and -alpha (IFNγ and IFNα) in Epi-*Dgcr8*^-/-^ caput epididymis and an RNA helicase important for spermatogenesis (DDX25) in GC-*Dgcr8*^-/-^ caput epididymis (Fig. 4F-G) (Tsai-Morris et al., 2004).

**Figure 4:**
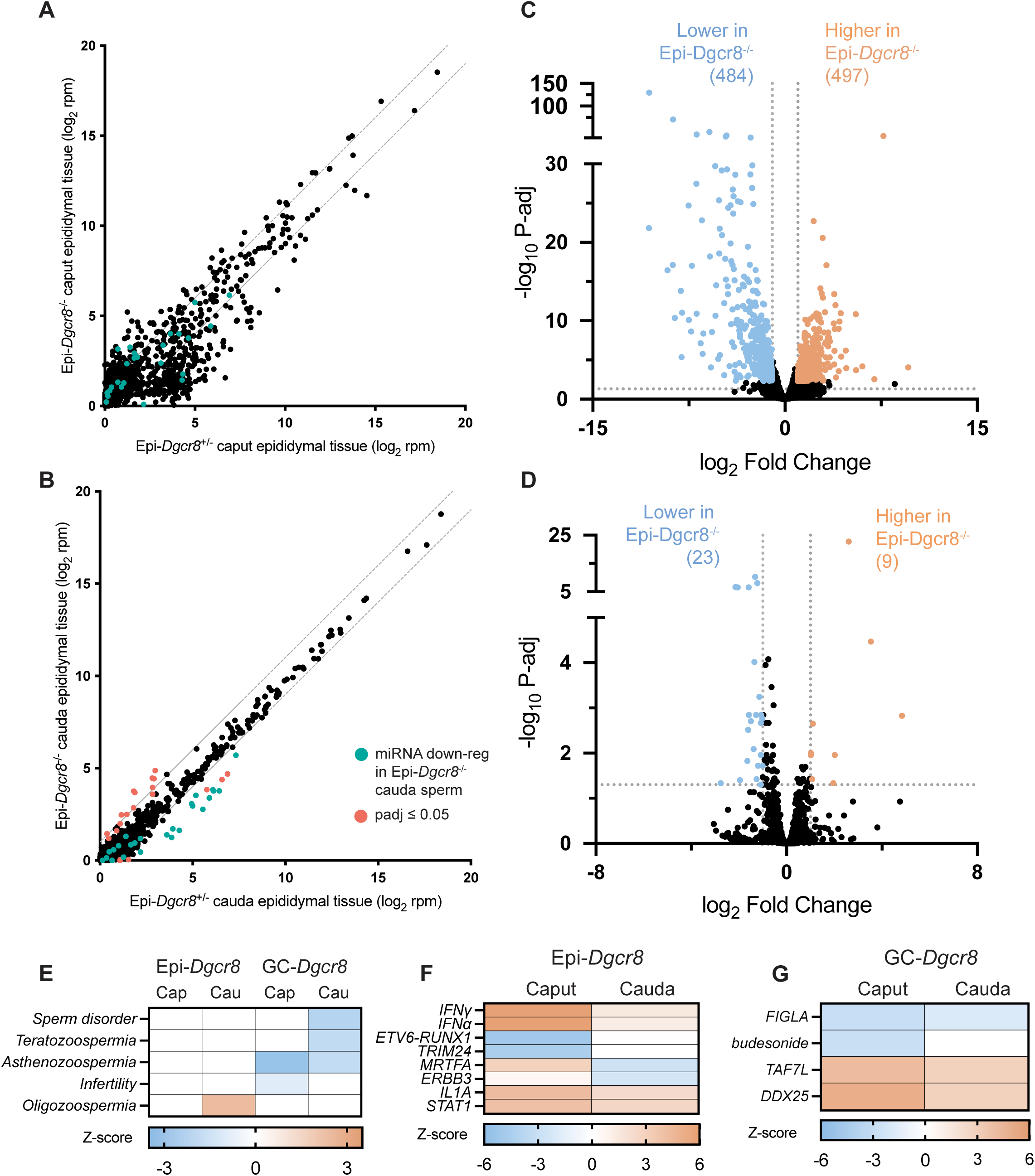
Loss of *Dgcr8* in germ cells and the proximal epididymis impacts miRNA and mRNA abundance in the male reproductive tract. (A-B) Scatter plot of microRNA (miRNA) levels in (A) caput and (B) cauda epididymal tissue of Epi-*Dgcr8*^+/-^ (x-axis) versus Epi-*Dgcr8*^-/-^ (y-axis). Data is normalized to total genome mapping reads (reads per million; rpm). Orange dots denote significantly altered miRNAs and teal dots indicate the 25 miRNAs reduced in Epi-*Dgcr8*^-/-^ cauda sperm. (C-D) Volcano plot depicting the log_2_ fold change and log_10_ adjusted *P*-value of mRNA-seq data from (C) caput and (D) cauda epididymis isolated from Epi-Dgcr8^-/-^ compared to Epi-Dgcr8^+/-^ male mice. Significance cutoffs of fold change > 2 and *P*-adj ≤ 0.05. (E) Heatmaps depict significantly impacted reproductive pathways identified in epididymal tissue samples from Ingenuity Pathway Analysis (IPA) of differentially expressed genes. Heatmap illustrating the upstream regulators as determined by IPA of differentially expressed genes in (F) Epi-*Dgcr8* caput and cauda epididymis and (G) GC-*Dgcr8* caput and cauda epididymis. A Z-score of ≥ 2 or ≤ -2 indicates the predicted activation or inhibition, respectively, of the upstream regulator.

### Epididymal acquired microRNAs influence preimplantation embryonic gene expression

To examine whether altered levels of miRNAs in sperm influence gene expression post-fertilization in the early embryo we first generated embryos fertilized by GC-*Dgcr8*^-/-^ testicular or cauda sperm, as well as the accompanying heterozygous sibling controls via ICSI. If sperm miRNAs regulate early embryo gene expression, due to the restoration of miRNA levels in GC-*Dgcr8*^-/-^ sperm following epididymal transit, we predicted that embryos fertilized by GC-*Dgcr8*^-/-^ testicular sperm would display gene expression changes that were not bestowed in embryos fertilized by GC-*Dgcr8*^-/-^ cauda sperm. Embryos were collected at the 4-cell and morula stage of development for assessment of gene expression by single embryo RNA-seq. Assessment of gene expression in embryos fertilized by GC-*Dgcr8*^-/-^ testicular sperm revealed dysregulated expression of 113 and 134 genes in 4-cell and morula stage embryos, respectively (*P*-value ≤ 0.01; Fig. S4A, 5A). Specifically, at the morula stage, expression of 101 of these genes was increased and 33 were decreased in GC-*Dgcr8*^-/-^ testicular sperm fertilized embryos compared to controls. In contrary, RNA-seq of GC-*Dgcr8*^-/-^ cauda sperm fertilized embryos revealed 40 down-regulated genes and no up-regulated genes at the 4-cell stage, while analysis of GC-*Dgcr8*^-/-^ cauda sperm fertilized morula embryos detected 15 down-regulated and 31 up-regulated genes compared to control embryos (grey dots Fig. S4B, 5B). Interestingly, only 7 DEGs were shared (up-regulated) in morula stage embryos when fertilized by either testicular or cauda GC-*Dgcr8*^-/-^ sperm compared to control sperm, thus aligning with the restoration of GC-*Dgcr8*^-/-^ cauda sperm miRNA levels during epididymal transit (Fig. 5C). Further, in examining the 134 differentially expressed genes in GC-*Dgcr8*^-/-^ testicular sperm fertilized morula embryos in GC-*Dgcr8*^-/-^ cauda sperm embryos, 93% of these genes (124 genes) were detected at levels similar to controls (orange and blue dots, Fig. 5B), a finding that was also mirrored in 4-cell embryo (Fig. S4B).

**Figure 5:**
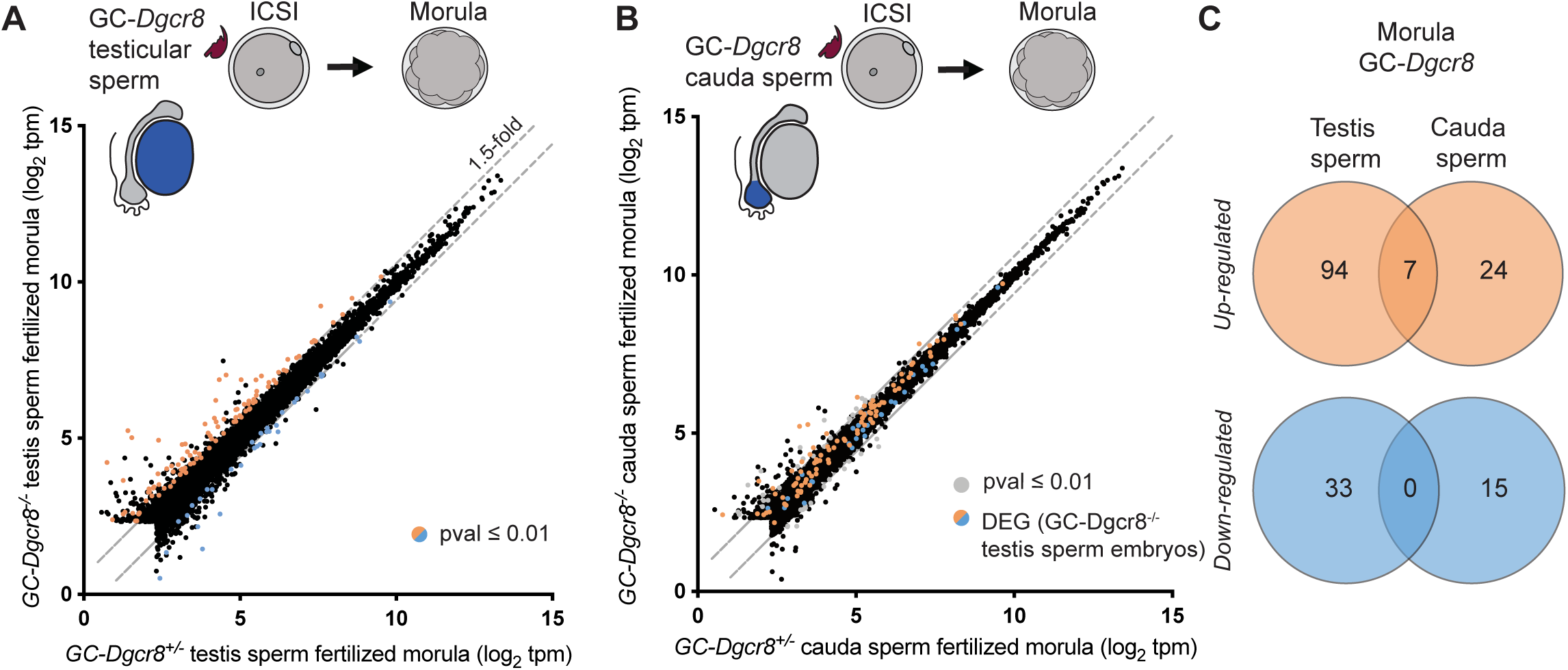
Gene expression analysis of germ cell *Dgcr8* knockout sperm derived preimplantation embryos. (A-B) Scatter plot of gene expression (transcript per million; tpm) in morula stage embryos fertilized by control sperm (x-axis, log_2_) versus GC-*Dgcr8*^-/-^ sperm isolated from (A) the testis and (B) cauda epididymis. Here, colored dots in both graphs represent the genes up-regulated (orange) and down-regulated (blue) in GC-*Dgcr8*^-/-^ testicular sperm fertilized embryo compared to control embryos. Data are shown for all genes with a mean expression of at least 5 tpm in one of the groups in the comparison. (C) Venn diagrams depicting the number of significantly up- and down-regulated genes shared between morula embryos fertilized by GC-*Dgcr8*^-/-^ testicular and cauda sperm.

Next, we sought to determine the influence of epididymal acquired sperm miRNAs on gene expression in the early embryo by fertilizing eggs via ICSI with Epi-*Dgcr8*^-/-^ sperm compared to that of heterozygous sibling controls. Transcriptomic analysis revealed 49 and 213 DEGs, in the 4-cell and morula stage embryo respectively (Fig. S4C, 6A, D). In the morula stage embryo, the majority of DEGs (184 genes) displayed increased expression in Epi-*Dgcr8*^-/-^ fertilized embryos (Fig. 6A). The observed gene expression regulation could be a result of aberrant sperm miRNA expression or additional impacts resulting from the loss of miRNA in the early epididymis independent of the miRNAs present in sperm. Therefore, to determine the extent of regulation attributed to sperm miRNA levels, we supplemented embryos with epididymal miRNAs (purified from cauda epididymosomes from wildtype mice) via microinjection. The supplementation of epididymal miRNAs to Epi-*Dgcr8*^-/-^ fertilized embryos led to a remarkable rescue of gene dysregulation in the early embryo (Fig. 6B). Specifically, of the genes altered in Epi-*Dgcr8*^-/-^ fertilized morula embryos, 65% (138 of 213 genes) returned to control levels in embryos that were subsequently microinjected with epididymal miRNAs, a phenomenon that was also evident at the 4-cell stage (Fig. 6C, S4D). Of note, several genes with highly significant (*P*<0.001) differential expression in Epi-*Dgcr8*^-/-^ embryos, including *Odc1, Srsf9* and *Hnrnpab* (Fig. 6D, S4F) are those that also display reduced expression in embryos fertilized by immature caput sperm, which have yet to acquire epididymal miRNAs (Conine et al., 2018; Sharma et al., 2018). Indeed, the expression of this subset of genes is restored when Epi-*Dgcr8*^-/-^ embryos are microinjected with epididymal miRNAs (Fig. 6E, S4E). Several pathways prominent among the differentially expressed genes in Epi-*Dgcr8*^-/-^ embryos when compared to control embryos were equally impacted when Epi-*Dgcr8*^-/-^ embryos were compared to miRNA-supplemented Epi-*Dgcr8*^-/-^ embryos, highlighting the pathways specifically influenced by epididymal miRNAs (Fig 6F). Pathways of note included IL-8 and GNRH signaling, which have both been implicated in preimplantation embryo development (Guzeloglu-Kayisli et al., 2009; Vilotić et al., 2022).

**Figure 6:**
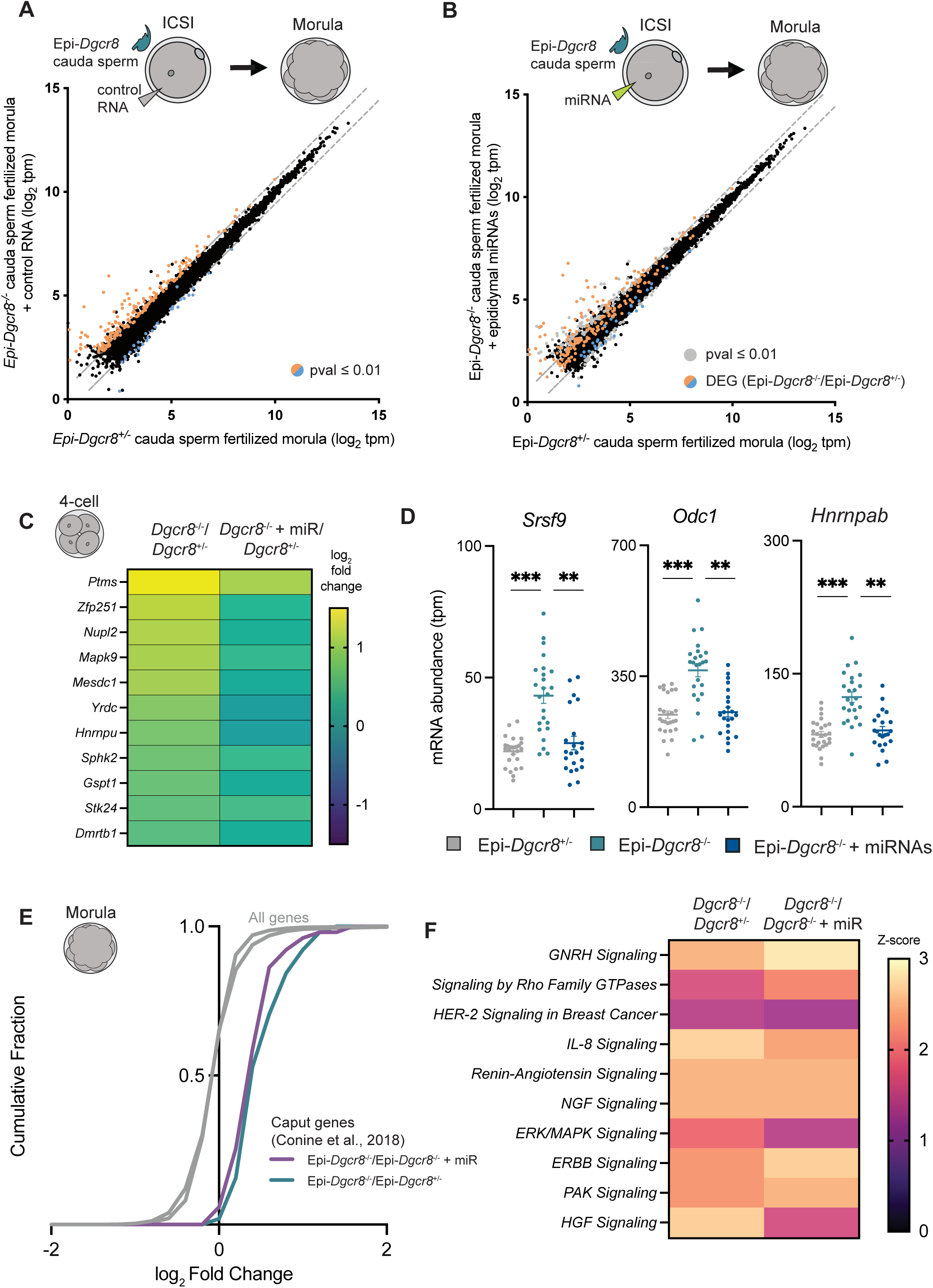
Epididymal-acquired miRNAs regulate preimplantation embryo gene expression. (A-B) Scatter plots of morula embryo gene expression (transcript per million; tpm) for control sperm fertilized (x-axis, log_2_) versus embryos fertilized by Epi-*Dgcr8*^-/-^ cauda sperm (y-axis, log_2_) then subsequently injected with (A) control RNA or (B) purified miRNAs from cauda epididymosomes. Here, colored dots in both graphs represent the genes up-regulated (orange) or down-regulated (blue) in embryos fertilized by Epi-*Dgcr8*^-/-^ sperm compared to control embryos. Gray dots indicate differentially expressed genes (*P* ≤ 0.01 and fold change 1.5). Data are shown for all genes with a mean expression of at least 5 tpm in one of the groups in the comparison. (C) Heatmap depicting the log_2_ fold change of the 11 genes significantly up-regulated in Epi-*Dgcr8*^-/-^ fertilized 4-cell embryos compared to control fertilized embryos. (D) mRNA abundance of individual morula embryos for three selected genes (*Srsf9*, *Odc1* and *Hnrnpab*). Horizontal bars represent average tpm. (E) Cumulative distribution function plot showing log_2_ fold change of all genes (grey) and a select group of genes that are up-regulated in embryos fertilized by caput sperm compared to cauda sperm (Conine et al., 2018) in Epi-*Dgcr8*^-/-^/Epi-*Dgcr8*^+/-^ (teal) and Epi-*Dgcr8*^-/-^/Epi-*Dgcr8*^-/-^ with miRNA supplementation (purple). (F) Top pathways identified by ingenuity pathway analysis (IPA) as activated (Z-score ≥ 2) or inhibited (Z-score ≤ -2) in Epi-*Dgcr8*^-/-^ embryos compared to control or miRNA injected Epi-*Dgcr8*^-/-^ embryos but not affected when control morula stage embryos were compared to miRNA injected Epi-*Dgcr8*^-/-^ embryos.

## DISCUSSION

Due to the inactive transcriptional state of mature sperm, the presence of RNA within sperm was originally thought to be mere remnants of testicular development (Amanai et al., 2006). However, over the past decade not only has the RNA cargo within sperm been assigned new functionalities, but the sperm RNA payload has proven to be dynamic, both physiologically during sperm maturation and in response to the paternal environment (Sharma, 2019; Trigg et al., 2019). In fact, the post-fertilization contributions of sperm small RNAs to offspring have been described most often in paternal intergenerational environmental effect studies, where numerous different exposures of males to stressors such as toxicants, and alteration in diet have been shown to transmit non-genetically inherited phenotypes to offspring. These phenotypes include altered metabolism, changes in behavior, and differential susceptibility to subsequent exposures (Chen et al., 2016; Rodgers et al., 2013; Wang et al., 2021), with the carrier of environmentally regulated non-genetic information causally linked to sperm small RNAs on numerous occasions, including both miRNAs and tRFs (Chan et al., 2020; Chen et al., 2016; Gapp et al., 2014; Rodgers et al., 2015). Additionally, embryos generated by sperm depleted of germline miRNAs and endo-siRNAs display altered preimplantation development (Yuan et al., 2016). Thus, in advancing our understanding of the contribution of sperm RNA to the developing embryo, it is critical to discern the source of sperm small RNAs. Here, we provide compelling evidence in support of the epididymis as a source of functional sperm miRNAs.

Sperm transit through the epididymis takes approximately 5-10 days in mammals (França et al., 2005; Robaire et al., 2006). Therefore, studies of acute paternal exposures encompassing this time period demonstrate that changes in the sperm small RNA profile, such as acrylamide exposure in mice (Trigg et al., 2021) and a high sugar diet in humans (Nätt et al., 2019) are the result of the dynamic response of the epididymis in directing sperm molecular changes under altered environmental conditions. While these studies and others suggest that small RNAs can be transferred from the epididymis to sperm, it has yet to be definitively demonstrated *in vivo*. Using a genetic approach, we identify the epididymal epithelium as a source of sperm miRNAs. Further, we report the restoration of miRNAs that were reduced in testicular sperm by virtue of *Dgcr8* loss in the germline, during epididymal transit, supporting the hypothesis that the epididymis is a site of sperm small RNA regulation following paternal environmental exposure. Albeit many of the miRNAs reduced in GC-*Dgcr8*^-/-^ testicular sperm encompass the same subset of miRNAs gained during epididymal transit (Fig. 3E). Thus, whether the epididymis is capable of sensing and restoring a subset of different miRNAs that are altered following environmental exposure and not already destined to be delivered during epididymal transit, remains to be determined. Indeed, some of the 25 epididymal acquired miRNAs identified in this study, including miR-880 and miR-34c, have been shown to be significantly altered in sperm of exposed fathers, including stress and altered diet (de Castro Barbosa et al., 2016; Dickson et al., 2018). As epididymal sperm are incapable of exerting a transcriptional response, such modulation to the sperm miRNA payload must be dependent on factors already present in sperm or driven by a mechanism of intercellular small RNA delivery. Owing to the established role of epididymosomes in the transfer of fertility modulating proteins (Martin-DeLeon, 2015), and the evidence presented here of the ability of miRNAs purified from epididymosomes to rescue embryonic gene expression changes that result from sperm lacking epididymal miRNAs, these vesicles are the leading candidate in orchestrating small RNA modulation to epididymal sperm. However, additional potential mechanisms have been postulated. These include the delivery of small RNAs via association with RNA-binding proteins (RBPs), through nanotubes and/or the shuffling between sperm and their cytoplasmic droplet (Battistone et al., 2019; Wang et al., 2023; Wang et al., 2010). At present, while plausible, there has been little investigation into the part that RBPs play in small RNA delivery in the epididymis. Moreover, nanotubes are reported to extend from clear cells of the epithelium, while the expression of *iCre* recombinase in this study is restricted to the principal cells (Björkgren et al., 2012); therefore, any contribution of small RNAs from the clear cells should not be affected. Further, the cytoplasmic droplet of mouse sperm harbors comparatively low amounts of miRNAs (∼2% of the total small RNA pool) (Wang et al., 2023). Hence, together with our current findings that the loss of *Dgcr8* in the epididymal epithelium results in populations of sperm with reduced levels of miRNAs (Fig. 3D), strongly supports the notion of an epididymal origin of sperm miRNAs.

Of equal importance to understanding where sperm acquire miRNAs is the identity of the specific miRNAs that are gained and whether these acquired miRNAs function post-fertilization to regulate early embryogenesis. Of the 25 miRNAs significantly reduced in Epi-*Dgcr8*^-/-^ sperm, 15 belong to the miR-880 X-chromosome cluster, as well as 3 from the miR-34/449 family, which are well-reported and reproducibly abundant miRNAs in mouse cauda sperm (Nixon et al., 2015; Sharma et al., 2018). In line with their reduction in Epi-*Dgcr8*^-/-^ sperm, these miRNAs have been reported to be enriched in cauda sperm and epididymis when compared to the caput (Belleannée et al., 2012; Nixon et al., 2015; Sharma et al., 2016; Sharma et al., 2018). The miR-880 X-chromosome cluster miRNAs, collectively referred to as Fx-miRs (Fragile-X miRs; also known as ‘spermiR’ and ‘XmiR’) and found at syntenic regions of the X-chromosome (between two conserved protein-coding genes, *Slitrk2* and *Fmr1*) are equally abundant in the testis and mature sperm in all mammals, including humans (Ota et al., 2019; Ramaiah et al., 2019; Zhang et al., 2019a). Moreover, reduced levels of these miRNAs are reported in sperm of human patients presenting with unexplained asthenozoospermia (Qing et al., 2017). Likewise, the altered abundance of miR-34/449 miRNAs in sperm has also been linked with male infertility, in mice and humans (Abu-Halima et al., 2013; Pantos et al., 2021). Thus, the abundance of these miRNAs in populations of mature sperm appears to be a critical indicator of sperm functionality and competency, which our findings also support.

While the expression of these miRNAs is undoubtedly important for spermatogenesis, we report here that among other miRNAs, the delivery of Fx-miRs and miR-34/449 via cauda sperm to the egg upon fertilization is important for the regulation of post-fertilization embryonic gene expression. Interestingly, despite the reduced expression of over 100 miRNAs in GC-*Dgcr8*^-/-^ testicular sperm, the array of altered genes in resultant embryos is far fewer than embryos fertilized by Epi-*Dgcr8*^-/-^ sperm. The miRNAs reduced in GC-*Dgcr8*^-/-^ testicular sperm overlap with Epi-*Dgcr8*^-/-^ cauda sperm, including a significant reduction in Fx-miRs. However, relatively, testicular sperm carry more copies of these miRNAs than cauda sperm (Sharma et al., 2018), and when *Dgcr8* is lost in the germline the levels of these miRNAs are subsequently reduced but remain at levels similar to control cauda sperm (Fig. S2F). Indeed, in examining the levels of these miRNAs in the populations of sperm used to generate embryos for RNA-seq it is evident that the sperm population harboring the lowest abundance is Epi-*Dgcr8*^-/-^ cauda sperm, which upon fertilization resulted in the greatest embryo gene expression changes. Further, these gene expression changes were rescued when miRNAs purified from cauda epididymosomes, which are abundant in Fx-miRs, are injected into Epi-*Dgcr8*^-/-^ embryos (Reilly et al., 2016). Therefore, this specific subset of miRNAs regulates gene expression upon delivery to the egg. Future studies to determine whether gene expression changes are a result of the collective down regulation of these miRNAs or driven primarily by only a few individual miRNAs warrants investigation. Notably, the ablation of one Fx-miR family, the miR-465 family in mice does not impair male fertility but leads to skewed sex ratios in resulting offspring due to dysfunction in the placentas of females (Wang et al., 2022).

Our findings that tissue specific epididymal ablation of the miRNA pathway produces sperm depleted of a subset of miRNAs, paired with the discovery that germ cell ablation of miRNAs recovers during epididymal transit, provides decisive genetic evidence of the soma-to-germline transfer of RNA in a mammalian system. This mechanism provides the environment a means by which it can modulate inherited regulatory information in sperm, via small RNAs and influence the phenotype of offspring (Chan et al., 2020). An open question remains how the epididymis senses and responds to the environment. Nevertheless, numerous studies have identified environmentally induced alterations in the epididymal soma that could conceivably contribute to the delivery of an altered small RNA payload to sperm, including modulation to transcription factors and RNA methyltransferases (Rompala et al., 2018; Trigg et al., 2021; Zhang et al., 2018). Moreover, a recent study has linked the microbiome and paternal T cells with the transmission of transgenerational non-genetically inherited phenotypes to offspring, opening the possibility of the epididymis relaying information in the form of RNAs loaded into sperm sensed by the immune system (Harris et al., 2023). Collectively, the findings of this study demonstrate that one function of the epididymis is to allow for molecular communication between the soma and germline, thereby breaking the “Weismann barrier”, by providing sperm with miRNAs which can influence embryonic development post-fertilization and non-genetically transmit phenotypes inherited by offspring.

## ACKNOWLEDGEMENTS

The authors gratefully acknowledge Rachel Bartlett, Liana Savarirayan and Madeline Lamonica for mouse husbandry and members of the Conine Lab for critical reading of the manuscript. We also thank the University of Pennsylvania School of Veterinary Medicine Comparative Pathology Core for histology assessment. N.A.T is the recipient of a Lalor Foundation Postdoctoral Fellowship (2022, 2023). C.C.C is the recipient of a Pew Biomedical Scholars Award.

## FIGURE LEGENDS

**Supplemental Figure 1:**
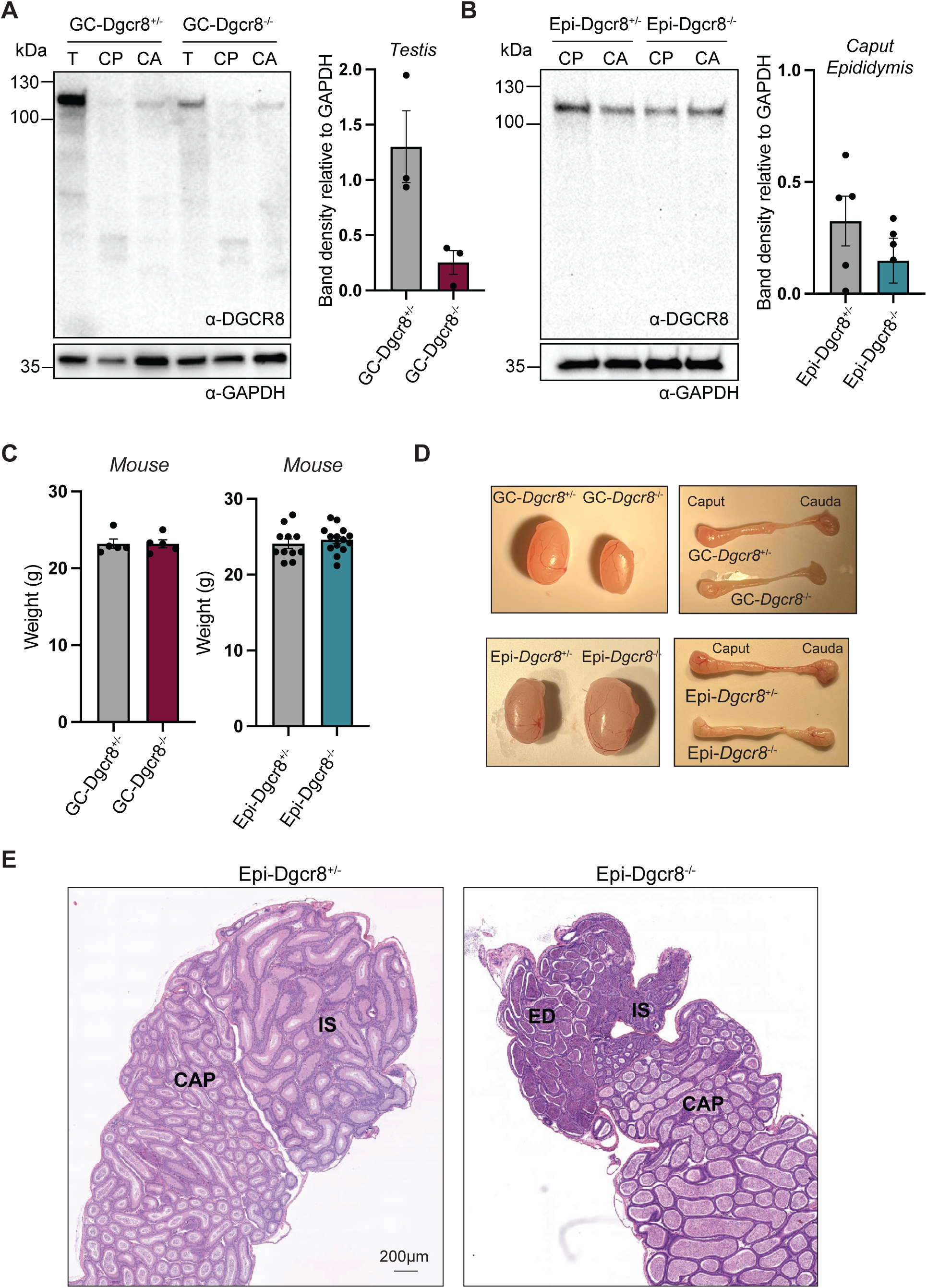
Conditional knockout of *Dgcr8* in the male reproductive tract. (A-B) Immunoblot analysis of DGCR8 protein expression in testis (T), caput (CP) and cauda (CA) tissue lysate of (A) GC-*Dgcr8*^-/-^ and (B) Epi-*Dgcr8*^-/-^ and control adult male mice. Experiments were performed at least three times and representative images are shown. Immunoblots were stripped and probed for anti-GAPDH to demonstrate equal protein loading. The density of the dominant DGCR8 band was quantified for (A) GC-*Dgcr8*^-/-^ testis and (B) Epi-*Dgcr8*^-/-^ caput epididymis and plotted as mean ± SEM relative to GAPDH. C) Whole body weight of experimental mice was measured after sacrifice. Data is presented as mean ± SEM. D) Images of dissected testis and epididymis from GC-Dgcr8^-/-^ and Epi-Dgcr8^-/-^ and control male mice. E) H&E staining of the proximal epididymis of Epi-*Dgcr8*^+/-^ and Epi-*Dgcr8*^-/-^ male mice. IS – initial segment, CAP – caput epididymis, ED – efferent duct.

**Supplemental Figure 2:**
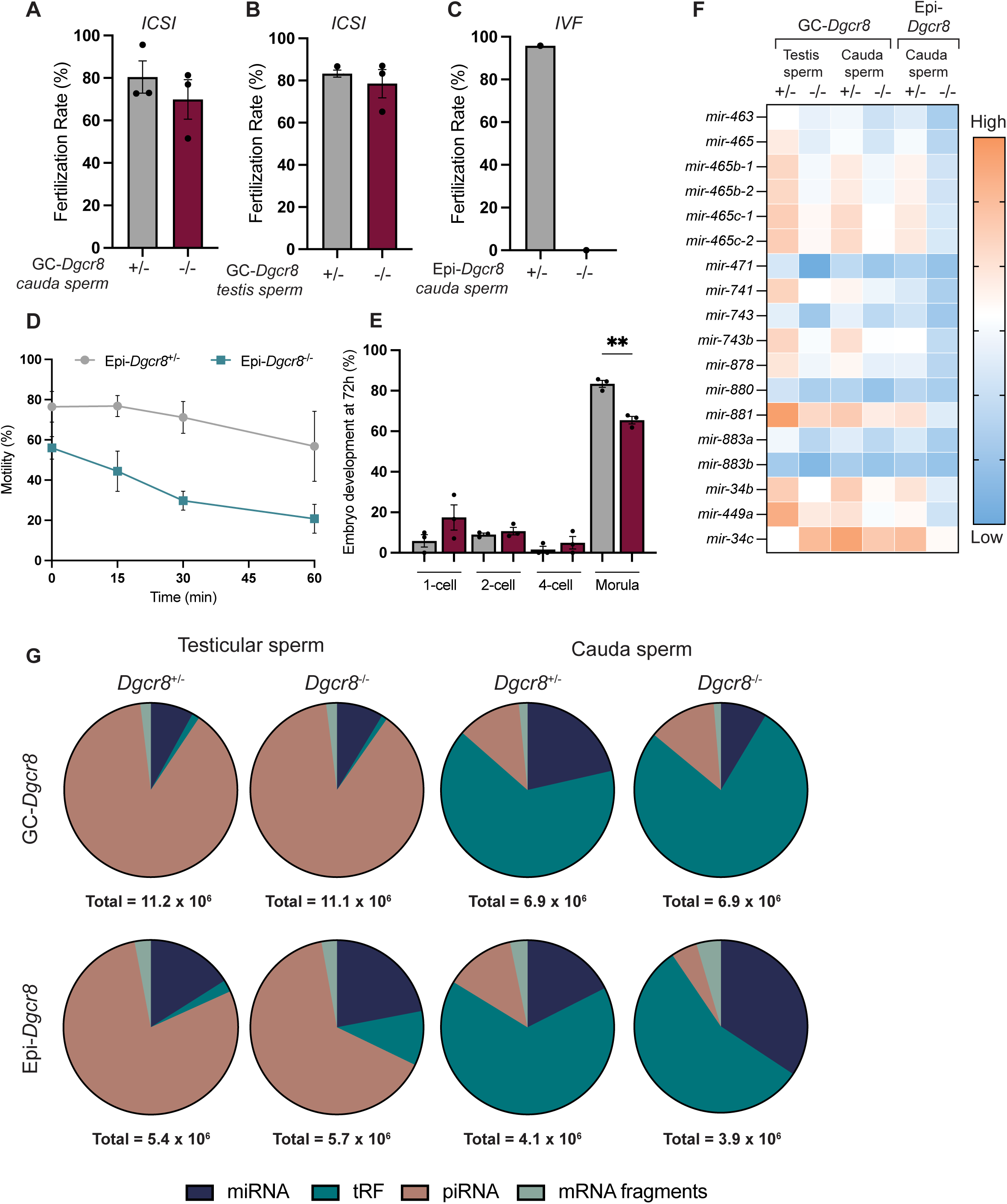
*Dgcr8* conditional knockout sperm functionality and small RNA-seq. (A-B) Fertilization rate of embryos fertilized by (A) GC-*Dgcr8*^-/-^ cauda sperm or (B) GC-*Dgcr8*^-/-^ testis sperm and the respective controls via intracytoplasmic sperm injection (ICSI). (C) Fertilization of eggs following in vitro fertilization (IVF) of Epi-*Dgcr8*^-/-^ and control cauda sperm (n=1). (D) Percent motility of Epi-*Dgcr8*^+/-^ and Epi-*Dgcr8*^-/-^ cauda sperm over 60 min *in vitro*. (E) 72 h development of embryos fertilized by GC-*Dgcr8*^-/-^ cauda sperm via ICSI compared to control sperm fertilized embryos. (F) Heat map of log_2_ expression count of epididymal acquired sperm miRNAs belonging to Fx-miR cluster and miR-34/449 family across all sperm populations. G) Contribution of each small RNA class to the global population of small RNA for testicular and cauda sperm isolated from control, GC-*Dgcr8*^-/-^, and Epi-*Dgcr8*^-/-^ mice. Pie charts depict average calculated across all replicates analyzed (n=4-8) Statistical significance is denoted as *P* ≤ 0.05 where asterisks represent levels of significance. ** = *P* ≤ 0.01.

**Supplementary Figure 3:**
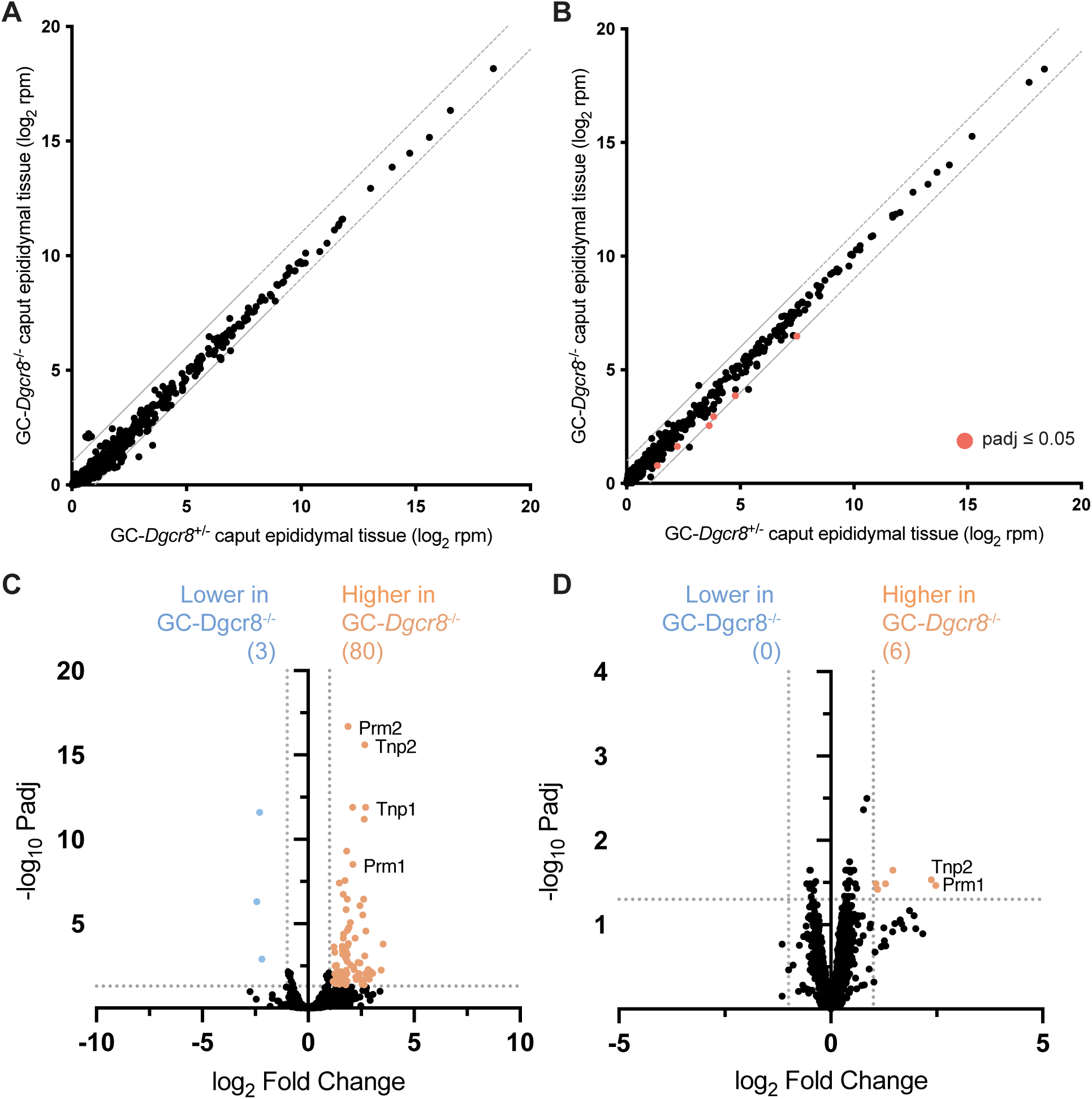
Epididymal miRNA and gene expression resulting from germ-cell *Dgcr8* ablation. (A-B) Scatter plots illustrating miRNA abundance in (A) caput and (B) cauda epididymis from GC-*Dgcr8*^-/-^ male mice (y-axis) compared to control mice (x-axis). (C-D) Volcano plot depicting gene expression changes in (C) caput and (D) cauda epididymis from GC-*Dgcr8*^-/-^ male mice compared to control mice. Colored dots indicate differentially expressed genes that satisfy the significance cutoff of *P*-adjusted value ≤ 0.05 and absolute fold-change ≥ 2 (dashed lines).

**Supplementary Figure 4:**
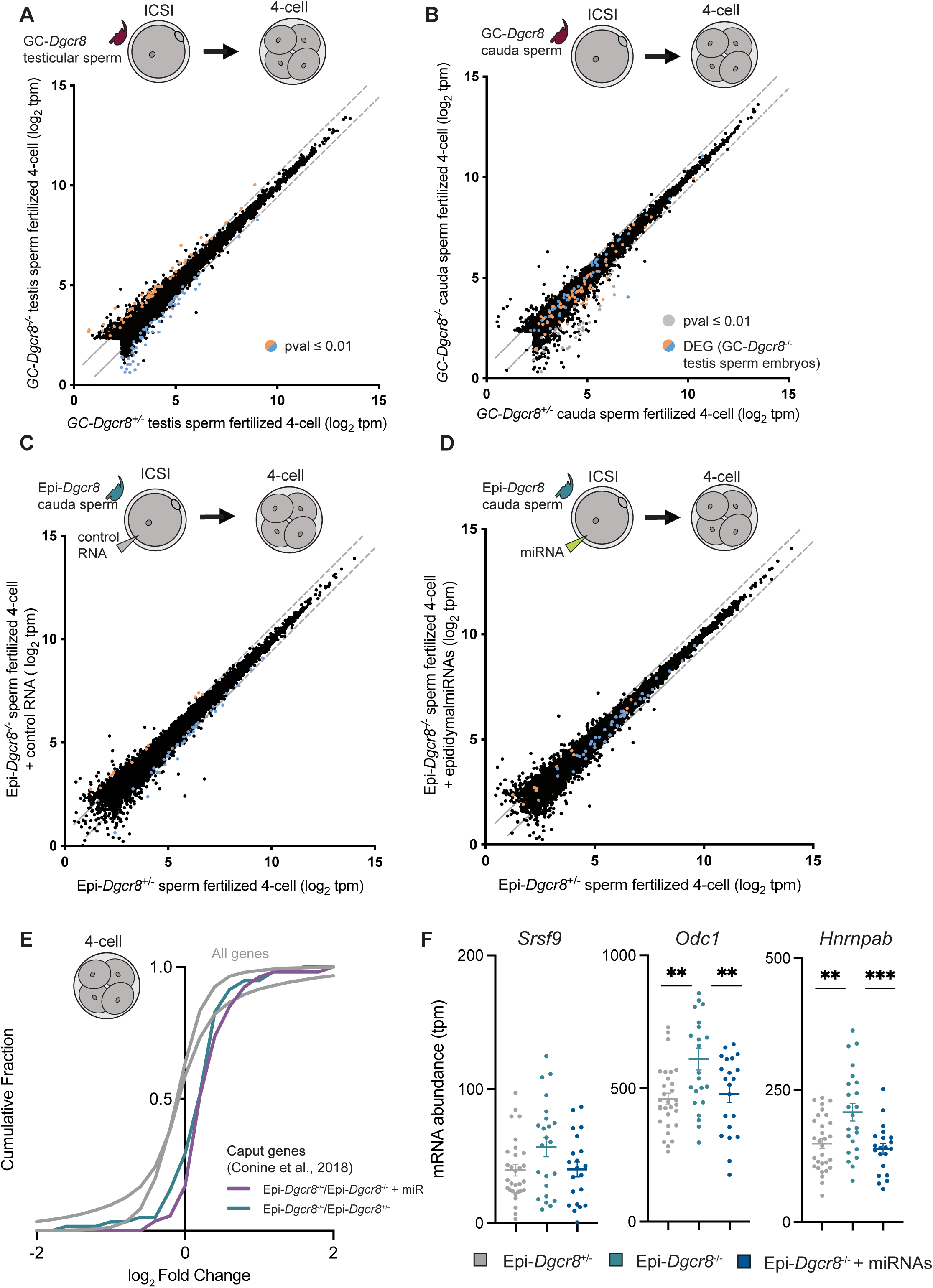
Embryonic gene expression of 4-cell embryos fertilized by *Dgcr8* knockout sperm. (A-B) Scatter plot of gene expression (tpm) of 4-cell embryos fertilized by control (x-axis) versus GC-*Dgcr8*^-/-^ A) testis and B) cauda sperm (y-axis). Colored dots in A and B indicate significantly altered genes (*P*-value ≤ 0.01) in embryos fertilized by GC-*Dgcr8*^-/-^ testis sperm compared to control testis sperm. (C-D) Gene expression (tpm) of 4-cell embryos fertilized by control cauda sperm (x-axis) versus 4-cell embryos fertilized by Epi-*Dgcr8*^-/-^ cauda sperm and subsequently injected with (C) control RNA or (D) miRNA purified from cauda epididymosomes. Here, colored dots in both graphs represent the genes up-(orange) or down-regulated (blue) in Epi-*Dgcr8*^-/-^ embryos compared to control embryos. Scatter plots depict data for genes with a mean expression of ≥ 5 tpm in at least one of the groups analyzed. E) Cumulative distribution function plot showing log_2_ fold change of all genes (gray) and a select group of genes that are up-regulated in embryos fertilized by caput sperm compared to cauda sperm (Conine et al., 2018) in Epi-*Dgcr8*^-/-^/Epi-*Dgcr8*^+/-^ (teal) and Epi-*Dgcr8*^-/-^/Epi-*Dgcr8*^-/-^ with miRNA supplementation (purple) 4-cell embryos. F) mRNA abundance of individual 4-cell embryos for three selected genes (*Srsf9*, *Odc1* and *Hnrnpab*). Horizontal bars represent average tpm.

